# Comparative morphology of silk-spinning systems in amphipods

**DOI:** 10.64898/2026.05.07.723571

**Authors:** Siena A. McKim, Thomas L. Turner

## Abstract

Silk glands have been found in two groups of amphipods: the Corophiida and the Ampeliscidae. The silk glands in Ampeliscidae, however, have yet to be examined in detail. Here we report, for the first time, the morphology and distribution of pereopodal glands in the Ampeliscidae, in non-thread producing Synopiidae, and in the Paragammaropsidae. In the Ampeliscidae we found two gland types distributed throughout all pereopods which have the ability to create threads. Pereopods three and four have additional silk extrusion morphology at the tip of the dactylus in which silk is transformed into semi-cylindrical threads used for building domiciles. Synopiid outgroup species have one of the gland types but lack silk extrusion morphology. Using ancestral state reconstruction analysis, we find that glands in the Synopiidae are likely ancestral and hypothesize that silk glands in Ampeliscidae are derived from these ancestral glands. Silk-spinning pereopods in the Paragammaropsidae had similarities with both Corophiida and Ampeliscidae but had distinctions. Ampeliscidae silk-spinning systems bear surprising resemblance to the Corophiida which presents one to reconsider the taxonomic placement of Ampeliscidae and the origins of silk-spinning in amphipods. This is the first comprehensive study on the glandular systems of Ampeliscidae, Synopiidae, and Paragammaropsidae using advanced microscopy, providing pertinent morphological data to the study of arthropod silk gland evolution and complex traits.

## INTRODUCTION

Silk is a proteinaceous material that is stored as a liquid stock and crystalized into a solid thread as it exits the body. The ability to spin a semi-cylindrical silk thread has allowed arthropods to interact with their environment in unique ways, contributing to their success worldwide (Mariano-Martins 2020). Silk-like proteins are found in many animal groups, but the ability to spin silk threads is known only in the arthropods (McDougal *et al*., 2016; Jung *et al*., 2021). Within this large phylum, however, silk production has evolved over 30 times (Craig 1997; Sutherland *et al*., 2010). The biology of silk production is most well-known in spiders and silk moths, which evolved silk production independently, but it is also found in several groups of crustaceans including tanaids, isopods, ostracods, and amphipods (Gorvett 1952; Wouters & De Grave, 1992; Kakui *et al*., 2021; McKim & Turner 2024). Marine silk has has likely been key to the success of several groups of amphipods, allowing thousands of species to create protective domiciles and produce feeding aids to catch food particles in the water column (Barnard 1969; Shillaker & Moore 1978; Dixon and Moore 1997)

Two main groups of amphipods are known to produce silk threads: Corophiida Leach, 1814 sensu Lowry & Myers, 2013 and Ampeliscidae Krøyer, 1842. The corophioids are a large infraorder within the suborder Senticaudata that includes species such as the giant kelp curler (*Sunamphitoe humerali*s Stimpson, 1864) and the European mud scud *(Corophium volutator* Pallas, 1766). Silk-spinning corophioids use a two-gland system in their pereopods three and four to spin marine silk (Kronenberger *et al*., 2012). Through histological staining of corophioid silk-spinning pereopods, previous studies have demonstrated that one gland type, often referred to as the proximal gland, produces a more acidic mucopolysaccharide or glycoproteins product while the other gland type, referred to as the distal gland, produces an acidic proteinaceous product (Kronenberger *et al*. 2012). Both gland types have been described as tegumental glands based on the crustacean gland literature (Talbot & Demers 1993) and more recently these glands have been called pseudotubular, due to their extreme elongation (Neretin *et al*., 2017). The proximal glands are found most densely in the basis but also found in the ischium and merus. The distal glands are most abundant in the merus and carpus. Historically, emphasis has been placed on the glands in the basis as they are more easily observable with the naked eye. Products from both glands appear to travel down the pereopod through central ducts, splitting into ductules in the dactylus, and then mixing before entering a chamber where they are stored before exiting the body (Kronenberger *et al*., 2012; Neretin 2016). The ductules and chamber appear to play a role in transforming the silk from a liquid feedstock to solid threads. This reservoir chamber is unique to corophioids and has yet to be described in any other arthropod silk-spinning systems (Neretin 2016). From corophioid morphology alone, it appears that amphipod silk requires mixing two products to produce a final spun silk. On other pereopods of corophioids, thread boring pores on the cuticle and secretory setae have been observed, suspected to produce mucous products contributing amorphous secretions to domicile construction (Mattson & Cedhagen 1989). More recent studies, however, lack investigation into these systems and little is known about these other glands.

Ampeliscidae is also a fairly large group that includes species like *Ampelisca abdita* Mills, 1964 and *Haploops tubicola* Liljeborg, 1856 and has been placed in an entirely different super order from the coriophids, the Amphilochidea Boeck, 1871 (Barnard & Karaman 1991). As a result, we expect that silk production evolved separately in these two groups of amphipods. In contrast to the corophioids, much less is known about the silk production system in ampeliscoids. Glands have been reported from the merus of pereopods three and four (Schellenberg 1925; Barnard & Karaman 1991; Cadien 2004) but it is unresolved how many gland types there are and how silk is transformed at the tip of the dactylus. Glands have also been found in other pereopods. Rigolet *et al*. (2011) stained the second pereopods of *Haploops nirae* with periodic acid Schiff–Alcian blue, which revealed large acid mucous glands in the propodus. These appear to secrete sticky products used to catch particles during suspension feeding (Rigolet *et al*., 2011). It is unclear if these glands are similar to those in pereopods three and four and how widespread pereopodal glands are. Determining the distribution and identity of pereopodal glands is critical in understanding the silk-spinning systems of ampeliscoids and the potential evolutionary origins of these glands.

An additional family, the Paragammaropsidae Myers & Lowry, 2003 have also been said to produce silk from the merus of pereopods three and four, but we are aware of no data on their morphology in the literature, including in the original species and family description (Myers and Lowry 2003). This family has been difficult to place taxonomically due to its conflicting morphology, so it is unclear if it is affiliated with the Ampeliscidae or the Corophiida or perhaps represents an additional origin of amphipod silk. It is currently within the superfamily Aetiopedesoidea Myers & Lowry, 2003, together with the Aetiopedesoidea. Glands were found within the merus of one species of Aetiopedesoidea (*Aetiopedes gracilis* Moore and Myers, 1988) similar to Ampeliscidae; however, Moore & Myers placed the entire superfamily within the Corophiida (Moore and Myers, 1988). Clearly, additional data on the silk-spinning morphology of these species would be valuable.

Understanding the comparative morphology of the Ampeliscidae, Corophiida, and Paragammaropsidae silk systems would shed light on the patterns and processes in the evolution of complex traits and the systematics of amphipods in particular. In terms of complex trait evolution, understanding the similarities and differences of convergently evolved traits at multiple levels of organization can reveal the underlying constraints that pattern organismal diversity. Incredibly, evidence points to more than 30 independent evolutions of silk production within arthropods (Craig 1997; Sutherland *et al*., 2010; McKim & Turner 2024). The repeated evolution of this trait within arthropods, but not outside this group, seems to imply that the evolution of silk spinning is not truly “independent”, but rather that there is some deep homology that preadapts arthropods to silk spinning. For example, the presence of an exoskeleton that glandular secretions must pass through to exit the body may facilitate the evolution of nozzle-like structures that can withstand the forces required to crystallize silk. It has been hypothesized in spiders and moths that silk ducts evolved from chitinous spines allowing for the development of silk products and silk extrusion morphology (Craig 1997; Davies *et al*., 2013). It could be that cuticular glands used to create the liquid feedstock are homologous across amphipods and preadapted lineages to evolve specialized silk-spinning systems.

Understanding the comparative morphology of the glands, ducts, pores and other parts of the silk-spinning machinery in these amphipod groups is a crucial step towards a better grasp of these patterns and processes governing the evolution of novel complex traits. Comparative morphology of these groups may also inform amphipod taxonomy and systematics, because the morphology of pereopodal glands has been used as evidence of shared ancestry (Myers and Lowry 2003; Lowry and Myers 2013). Based on the results of Moore & Myers (1988) and Myers and Lowry (2003), we hypothesize that the silk gland system of Paragammaropsidae should be more similar to Ampeliscidae, due to prominent glands in the merus of pereopods three and four. Further delineating how the Paragammaropsidae produce silk should indicate their affiliation within the Amphipoda.

To advance these aims, we have used histological staining and advanced microscopy and spectroscopy to discover the silk-spinning morphology and glands in seven ampeliscoid species, one paragammaropsid species, and three synopiid species. We examined the spun silk of the domiciles from *Ameplisca lobata* and *Paragammaropsis sp.* to connect the pore morphology to its products. To produce a hypothesis regarding the origin and evolution of silk-spinning system traits, we estimated a phylogeny for these species and performed ancestral state reconstruction. We describe these systems in detail for the first time and compare them with corophioid silk-spinning systems.

## MATERIALS AND METHODS

### Phylogenetic Estimation and Analyses

We used publicly available data from the cox1 locus (the only locus with sequences available for a large number of species) to estimate a phylogeny for the Ampeliscidae. Using the NCBI taxonomy browser and BLAST searches, we located ∼658 bp cox1 sequences from 15 taxa within Ampeliscidae, along with 7 closely related outgroup taxa from the superfamily Synopioidea and one more distantly related outgroup, an arctic mysid. See S1 Table for the accession numbers for all genes used. Unfortunately, molecular data from *Paragammaropsis sp*. was not available. All sequences were screened for contamination and taxonomic inconsistencies using BLASTn (Altschul *et al*., 1990). The dataset was inspected and edited in AliView v1.18.2, and sequences were aligned with MUSCLE v3.8.31. Portions of the alignment containing long stretches of ambiguous homology on the 3’ end were removed prior to phylogenetic inference.

Phylogenetic relationships were inferred using both Maximum Likelihood (ML) and Bayesian inferences. For the ML tree, we used IQ-TREE 2.1.4_beta (Nguyen *et al*., 2014). The best-fitting nucleotide substitution model for the alignment was selected with ModelFinder (Kalyaanamoorthy *et al*., 2017) as implemented in IQ-TREE using the Bayesian Information Criterion (BIC). The selected model TIM+F+R4 was then used to estimate the maximum likelihood tree. Node support was assessed using 100,000 ultrafast bootstrap replicates (UFBoot; Minh *et al*., 2013).

Phylogenetic relationships were inferred using Bayesian inference in MrBayes v3.2. To account for variation in substitution patterns among codon positions, the sequences were partitioned by codon position: first, second, and third positions and defined as separate character sets. A three-partition scheme was applied to model each codon position independently. For each partition, the general time-reversible (GTR) substitution model with gamma-distributed rate variation among sites (four discrete categories) was applied. Base frequencies were set to empirical, and substitution rate matrices, base frequencies, and gamma shape parameters were unlinked across partitions, allowing each codon position to have independent parameter estimates. Two independent Markov Chain Monte Carlo (MCMC) runs were performed, each with four chains (one cold, three heated) for 15 million generations, sampling every 500 generations. Convergence was assessed by monitoring the standard deviation of split frequencies, and a burn-in of 25% of sampled generations was discarded. Runs converged (final ASDSF = 0.000683; maximum split-frequency SD = 0.002545; average PSRF = 1.000; maximum PSRF = 1.001). All key parameters had ESS > 200. The remaining trees were summarized to obtain the majority-rule consensus tree with posterior probabilities as branch support values.

The bayesian phylogeny provided a backbone for marginal ancestral state reconstruction (ASR) of silk gland traits. Silk gland traits included the presence/absence of gland type one and the presence/absence of small terminal pores on the dactylus of pereopods three and four. Using rayDISC within the corHMM package in R, we tested for the best evolutionary model for each ASR analysis and in both cases, the equal rates model with a maddfitz rootprior was the best model based on the AIC and AICc scores. The marginal probability of character states at each node was then represented as pie charts on the tree. The R code and files for ASR are available on github at coding4amphipods/Ampeliscidae_publication.

### Morphological Sampling and Microscopy

We investigated the pereopodal glandular morphology of 11 species. For Ampeliscidae representatives we examined *Ampelisca lobata* Holmes, 1908, *A. romigi* Barnard, 1954, *A. pacifica* Holmes, 1908, *Haploops setosa* Boeck, 1871, *H. laevis* Hoek, 1882*, Byblis veleronis* Barnard, 1954, and *B. millsi* Dickinson, 1983. As an outgroup to the Ampeliscidae, we examined several species from the Synopiidea Dana, 1852: *Syrrhoe longifrons* Shoemaker, 1964, *Bruzelia tuberculata* Sars, 1883, and *Tiron biocellatus* Barnard, 1962. We examined one species of Paragammaropsidae, *Paragammaropsis sp* Ren & Huang, 1991. The *A. romigi, A. pacifica, B veleronis, B. millsi, H. laevis, H. setosa, B. tuberculata* and *T. biocellatus* specimens were obtained from the Natural History Museum of Los Angeles. *S. longifrons* was loaned from the Santa Barbara Museum of Natural History. Specimens of *Paragammaropsis sp.* were collected in Antarctica by Kevin Kocot’s lab during the ICY INVERTS 2020 expedition and loaned by the Alabama Museum of Natural History. *A. lobata* specimens were collected locally in Santa Barbara by SCUBA during the UCSB 2025 Bioblitz. See S2 Table for additional specimen details.

In total we examined 112 pereopods through compound microscopy, confocal laser microscopy, micro-computed tomography, and scanning electron microscopy. To examine pereopods with compound and confocal microscopes, we first removed pereopods two to seven between the coxae and basis. These whole legs were then stained using bromophenol blue following the protocol from Kronenberger *et al*. (2012) and Bonhag (1955) (see S3 Protocol for full stain recipe and protocol). Stained legs were whole mounted on glass slides with a coverslip and imaged using 4-100x magnification on the Olympus BX 51 (Olympus, Tokyo, Japan) light microscope with a Lumenera Infinity 3 (Teledyne, Ottawa, Canada) camera in the UCSB Neuroscience Research Institute Microscopy Facility. For topographically complex structures, we used manual stacking, taking a photo every 10-40 µm and then compiling images to create one stacked image using the z-project function in Fiji (Schindelin *et al*., 2012). For confocal images, we used the Leica SP8 (Leica Biosystems, Wetzlar, Germany) confocal scanning laser microscope in the UCSB NRI-MCDB microscope facility on non-stained, whole mounted legs. Using a 488 nm excitation laser and dry 10x magnification, we captured the autofluorescence of the sample at 1028 x 1028 resolution.

The external morphology of some specimens was examined using Scanning Electron Microscopy (SEM) with a FEI Quanta 400F Field emission source SEM (FEI Company, Hillsboro, OR, USA) in the UCSB Department of Earth Science. Whole specimens were placed on an aluminum SEM stub with an adhesive carbon substrate and then coated with Gold/ Palladium using a sputter coater. The SEM was operated using a 5kV accelerating voltage and a beam current of ∼0.5nA. Specimens were imaged by collecting the secondary electron signal using an Everhart-Thornley electron detector. Photomicrographs were typically acquired by integrating 32 frames at 1024x884 pixel resolution with 1us dwell time per pixel per frame.

The third pereopod in a *Pargammaropsis sp.* specimen was analyzed using micro-computed tomography (micro-CT) to resolve the distribution of glands within the pereopod. The pereopod was stained for two weeks in phosphotungstic acid (PTA) on an orbital shaker. The specimen was then transferred to a 0.5 mL eppendorf tube, filled with 70% ethanol, and placed on top of wool. The scan was taken with a source of 40kv voltage, 200uA current, 8w power imaging with pixel size of 1.317, optical magnification of 9.8 “10x” and with an exposure time of 3.5 seconds. A Xradia MicroXCT-200 (Carl Zeiss, Oberkochen, Germany) scanner was used by Sebastian Büsse (University of Greifswald, Evolutionary Biology Department). Two-dimensional projection images were reconstructed into tomograms and then stacked and segmented using 3D Slicer (Kikinis *et al*., 2014) and Biomedisa (Lösel *et al*., 2020). A final three-dimensional volume rendering was produced.

## RESULTS

To investigate the evolution of silk-spinning systems within the Ampeliscidae, we first inferred a phylogeny using publicly available data from the cox1 locus (Fig. 1). When the phylogeny was estimated with maximum likelihood (ML), bootstrap values within some genera, such as *Ampelsica* and *Haploops*, were lower than 80. We used Bayesian inference to better resolve these relationships despite limited sequence data. The relationships among species within ampeliscoid genera were sometimes poorly resolved, but the relationships between genera had high posterior probabilities. The topology of the inferred tree was consistent between the Bayesian and maximum likelihood approaches, except for *Syrrhoites*, which was monophyletic in the ML tree but not in the Bayesian tree. This phylogeny supports the monophyly of the Ampeliscidae clade, as found in Copilaş-Ciocianu *et al*. (2020), and the monophyly of each ampeliscoid genus. *Haploops* and *Ampelisca* are inferred as sisters, with *Byblis* being sister to them. Synopidea, however, was not found to be monophyletic.

**Figure 1.**
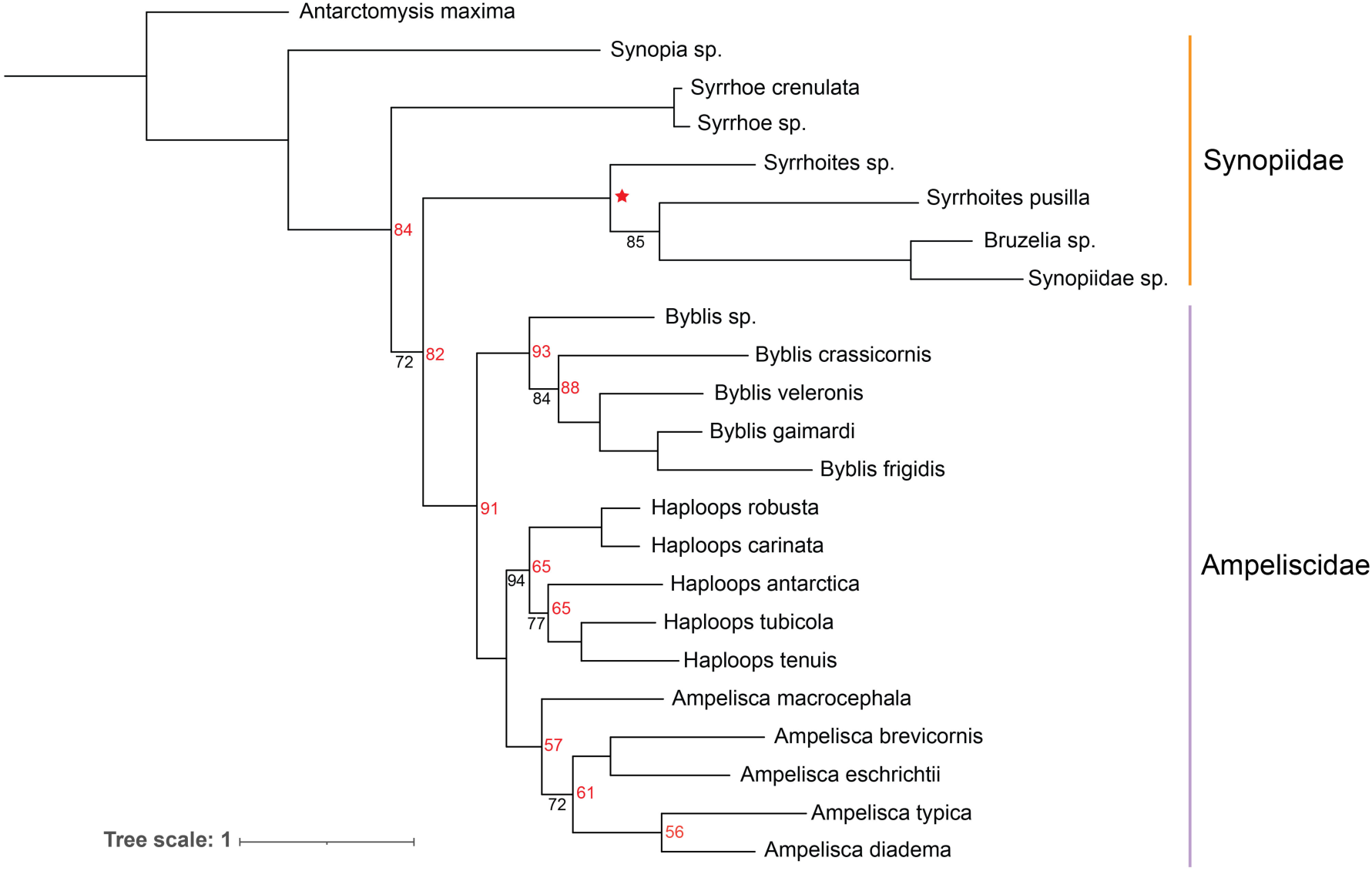
Estimated phylogeny. Phylogenies were estimated with both maximum likelihood and Baysian methods; the Baysian tree is shown. Nodes have bootstrap ≥ 95% and posterior ≥ 97%, except where indicated: red numbers indicate low bootstrap values, and black numbers indicate low posterior probabilities. The red star indicates the only node that wasn’t present in the maximum likelihood tree.

We then used microscopy to explore the pereopod gland morphology among these taxa. Replicate individuals were examined in seven species: three from the genus *Ampelsica,* and two species each from *Haploops* and *Byblis*. Glands were found in silk-spinning pereopods three and four across all ampeliscoid species examined (Fig. 2) consistent with previous reports (Mills 1967; Enequist 1949). We found that these glands, however, were not just present in the silk-spinning pereopods but were present throughout pereopods two through seven (Fig. 2B). Though previously reported only from the merus, glands within pereopods three and four occupied most of the hemocoel in the basis, ischium, merus and carpus; this was also seen in the other pereopods. Two distinct types of glands were observed when specimens were stained with bromophenol blue, which we term gland type one and gland type two. Gland type one had secretory cells dense with secretory vacuoles and stained intensely. Gland type two had secretory cells that stained weakly. Both gland types had stained accumulation sites, but these were darker in gland type two. Gland type two accumulation sites also tended to be larger. In all ampeliscoids, both gland types are distributed throughout all examined pereopods. Type one glands have accumulation sites with a lateral duct connected to a central duct that travels down the pereopod. In *Ampelisca* species, within a single individual, there was variation in type one glands in pereopods five to seven; some glands had secretory cells stained light blue or orange and others were stained dark blue (Fig. 2F). We term these gland types 1A and 1B, respectively.

**Figure 2.**
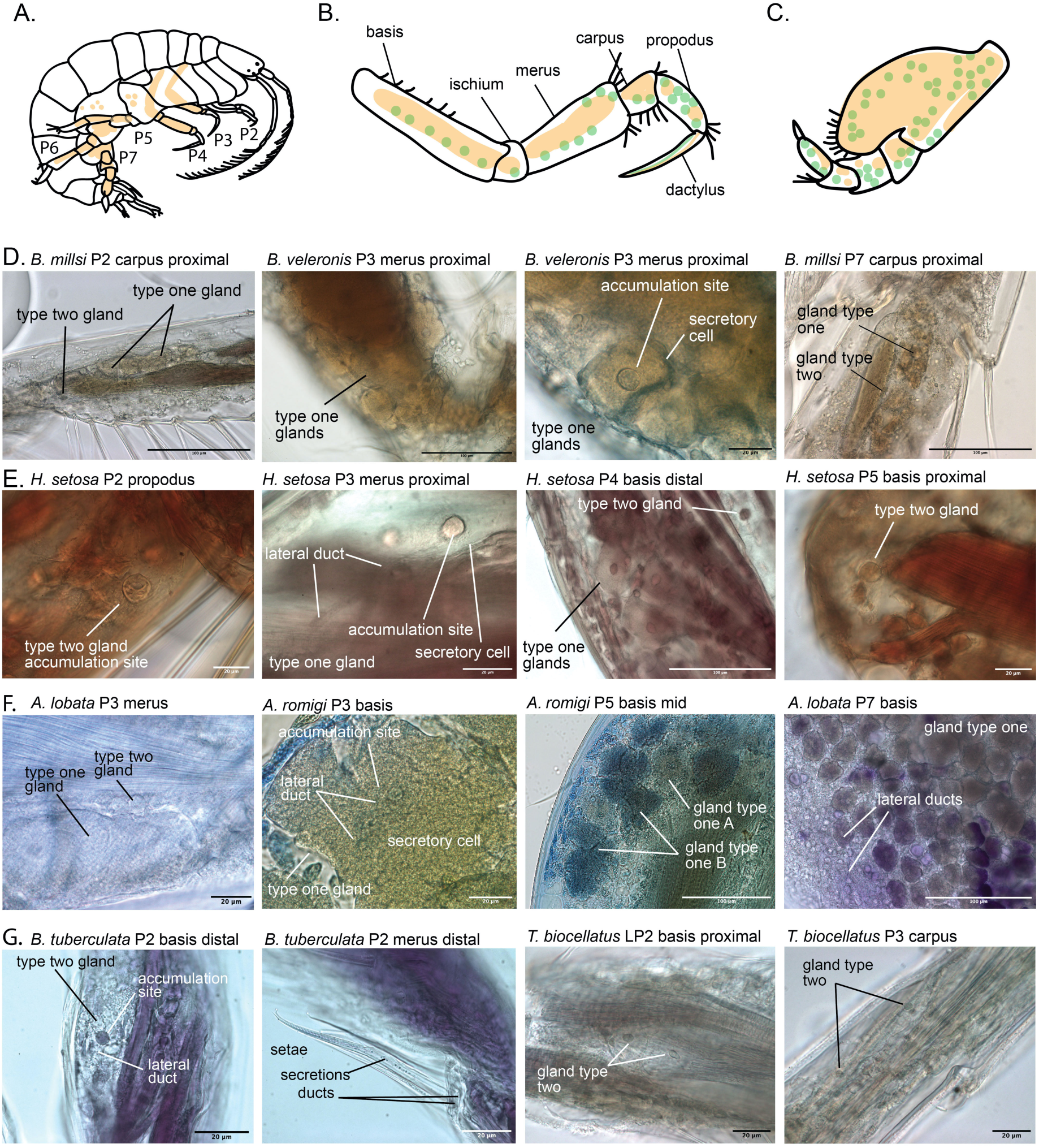
Gland morphology of ampeliscoids and synopiids. Illustration of a generalized ampeliscid with pereopod numbers labeled and orange indicating gland distribution (**A**). Generalized illustration of ampeliscoid silk-spinning pereopods three and four with type one glands (orange) and type two glands (green) (**B**). Generalized illustration of non-silk-spinning pereopod seven with gland type one and two (**C**). D-G Photographs of pereopod tissue stained with bromophenol blue; variation in colors is due to the varied preservation methods of the specimens. *Byblis* (**D**). *Haploops* (**E**). *Ameplisca* (**F)**. Synopiids *Bruzelia tuberculata* and *Tiron biocellatus* (**G**).

Using light microscopy and SEM, we examined pereopod three and four dactyls where, presumably, silk is transformed. Ampeliscoids had extremely elongated and narrow dactyls that the gland ducts traveled down to, creating specialized morphology to extruded silk on the distal tip (Fig. 3). We found no differences in the morphology of pereopod three and four. All genera had a channel running down the middle of the dactylus, however, the dactyl tip where secretions exited differed within each genus. *Ampelisca* (Fig. 3A) had a large proximal pore with another distal large pore alongside a group of small pores. The distal large pore appeared to have a small chamber proximal to the exit. In addition to two large pores, there were four to sixteen small pores on the distal end of the dactylus. The dactyl channel ran up the middle of the dactylus which connected to a groove with a seta (Fig. 3C). *Haploops* shared all these characters, but the chamber tended to be larger (Fig. 3B). *Byblis* differed the most in dactylar morphology. *Byblis* also had a chamber with a large pore but with two distal large pores instead of one. *Byblis* lacked terminal small pores. Our ASR analysis supports the origin of the small terminal pores to be in the most recent common ancestor of *Ampelisca* and *Haploops*, rather than in all ampeliscoids (Fig. 5B). *Byblis* also had a channel following the proximal setal groove but this channel, instead of being restricted to the ventral side, appeared to create a distinction between the dorsal and ventral planes of the dactylus, and connected to a second setal groove.

**Figure 3.**
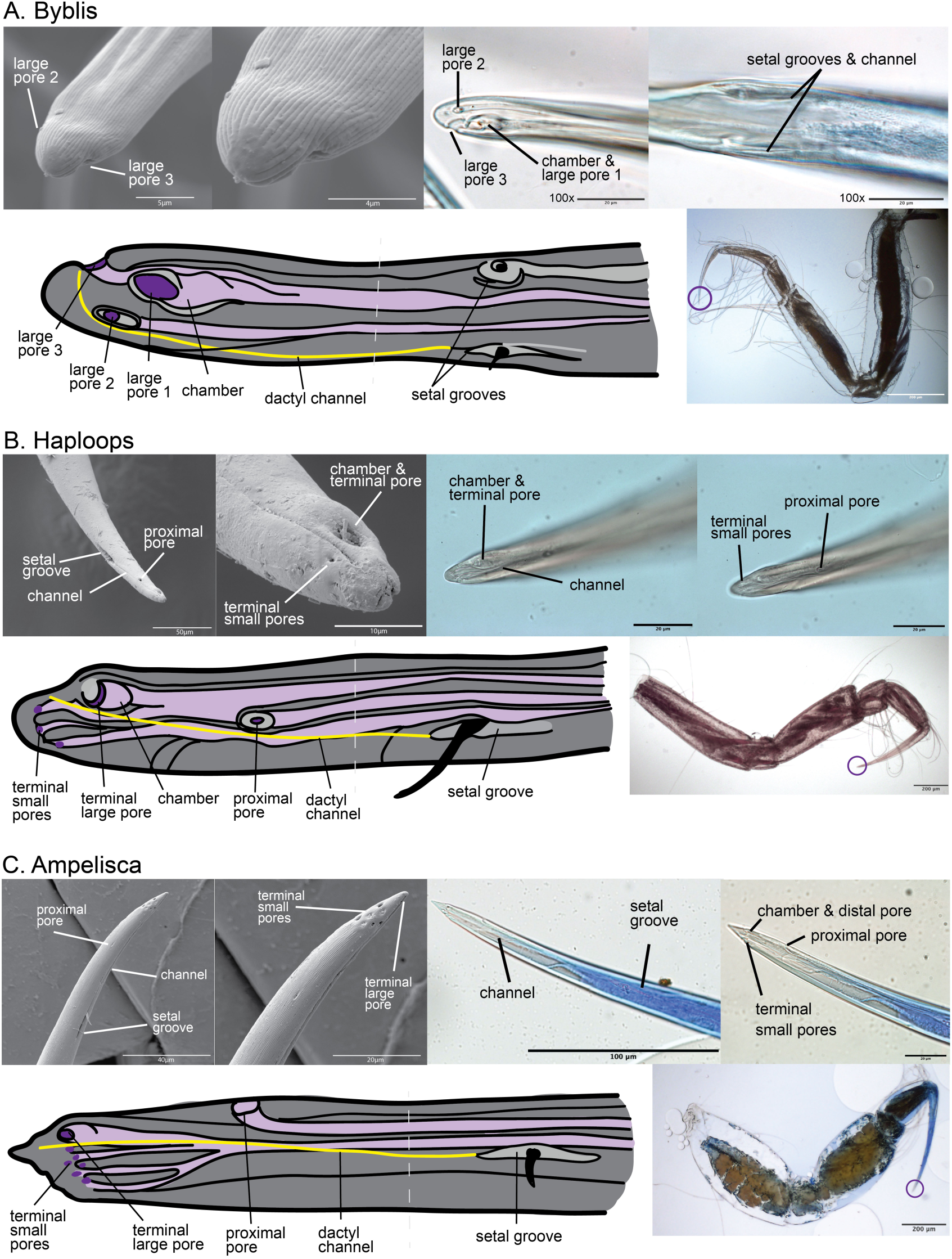
Silk-spinning dactylar morphology with images from SEM, light microscopy on bromophenol blue stained legs, and illustrated diagrams. Key characteristics include setal grooves and pores (light grey), secretion points from pores (dark purple), gland ducts (light purple), dactylar channel (yellow), and setae (black). *Byblis*; all images of *B. veleronis* (**A**). *Haploops*; all images of *H. setosa* (**B**)*. Ampelisca*; SEM of *A. lobata* and light microscope images from *A. romig*i (**C**).

The dactyl tip on pereopods three and four appears to be the main source of silk production. We found additional pores on the surface of the cuticle, however, which produced thin threads in two species: *A. lobata* (Fig. 6D & E) and *H. setosa* (Fig. 6G). In *A. lobata*, these threads do not appear to make the thick silk threads found in the domicile (Fig. 6A-C) but likely contribute to smaller thinner threads with an unknown function. On pereopod seven of *A. lobata* we found additional cuticle pores with threads (Fig. 6D) including on the posterior basis and merus in high density, where coincidentally we found a high density of both gland types. Both gland types were found in pereopod two, five, and six, however, we could not identify with confidence where the gland secretions exited the body. SEMs of the silk domicile from *A. lobata*, and silk on the specimen of *H. setosa*, show a variety of morphologies including thin threads, thick threads, and amorphous products, suggesting that different pores produce different products or altered product states.

To better understand the prevalence of pereopodal glands, and the origin of silk-spinning glands in ampeliscoids, we also examined the pereopods of closely related synpoiid species. Surprisingly, we found gland type two throughout all examine pereopods of synopiids. In *Bruzelia tuberculata*, we found gland type two prominently in pereopod two, where we observed secretory setae on the merus connected to ducts running up the pereopod to the glands (Fig. 2G). In other cases, such as *Tiron biocellatus,* it appears glands secrete to the cuticle surface (Fig. 2G). Pereopods three and four lacked the reservoir chamber, pores, and slits seen on the dactyls of silk-spinning ampeliscoids. See Table 1 for summary of pereopodal characters in all examined groups (entire comprehensive table available on request). Our findings of type two glands in outgroup synopiids, and the restriction of type one glands to ampeliscoids, prompted us to explore the ancestral state of glands using ASR analysis. Our analysis supports the origin of gland type one within the ampeliscoid common ancestor rather than in the common ancestor of synopiids, with gland type two as the ancestral state (Fig. 5).

**Table 1.** Pereopodal gland systems summary. 0 = character absent, 1 = gland type one, 2 = gland type two, 3 = both gland type one and two present, 4 = large other epicuticle glands found in *Paragammaropsis sp*.

| Taxon | glands<br>P2 | glands<br>P3 | glands<br>P4 | glands<br>P5 | glands<br>P6 | glands<br>P7 | silk-spinning<br>dactylus<br>P3 | silk-spinning<br>dactylus<br>P4 |
| --- | --- | --- | --- | --- | --- | --- | --- | --- |
| *Ampelisca* | 3 | 3 | 3 | 3 | 3 | 3 | yes | yes |
| *Haploops* | 3 | 3 | 3 | 3 | 3 | 3 | yes | yes |
| *Byblis* | 3 | 3 | 3 | 3 | 3 | 3 | yes | yes |
| *Paragammaropsis* | 4 | 1 & 4 | 1 & 4 | 4 | 2 & 4 | 4 | yes | yes |
| Synopiidae | 2 | 2 | 2 | 2 | 2 | 2 | no | no |

Lastly, we investigated the glandular systems of *Paragammaropsis sp*. which has an uncertain taxonomic placement. The glands of *Paragammaropsis* shared similarities with the glands of ampeliscoids and synopiids but had distinctions (Fig. 4). Glands were found within the merus and carpus of pereopods three and four. When stained with bromophenol blue, these glands had intensely stained secretory cells, similar to the type one glands of ampeliscoids (Fig. 4D). The accumulation sites of these glands were also massive (Fig. 4C & D). Glands that matched the type two glands of ampeliscoids were also seen, in the basis and merus of pereopod six, but these glands were fewer in number than in ampeliscoids. Large, oval, epicuticle glands with no obvious duct system were found throughout all pereopods; these lacked similarity to any glands seen in ampeliscoids (Fig. 4A). These glands lacked any obvious accumulation sites but appeared to be full of secretory vacuoles (Fig. 4C). *Paragammaropsis* had extremely elongated narrow dactyls on pereopod three and four, with distal secretory pores, similar to *Byblis*. On the dactyls, there were two setal grooves that extended mid dactylus to the tip creating a distinct channel between the dorsal and ventral regions of the tip (Fig. 4E-H). The dactylus tip, however, had many small pores, up to twenty, similar to *Ampelisca*. Similar to ampeliscoid silk, SEM of the *Paragammaropsis* domicile revealed thick threads, thin threads, and also sticky amorphous secretions (Fig. 6H & I).

**Figure 4.**
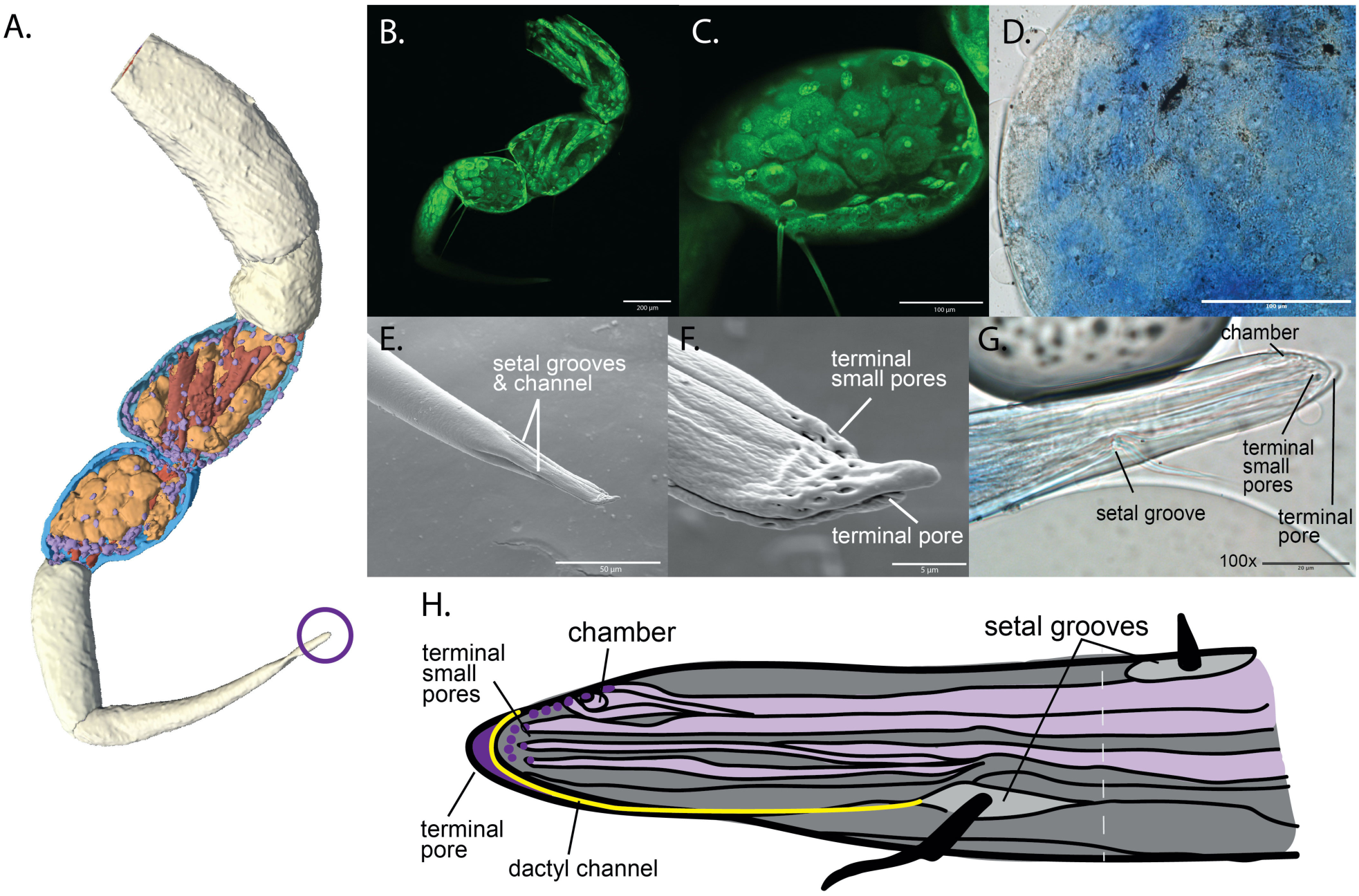
*Paragammaropsis sp*. pereopod silk-spinning morphology. Micro-CT volume reconstruction with merus and carpus sliced open highlighting type one silk glands (orange), epicuticle glands (purple), muscle (red), inside layer of cuticle (blue), and exterior cuticle (white) (**A**). Confocal image of autofluorescent pereopod four, basis, ischium, merus, carpus, and part of the propodus. Silk glands in the merus and carpus (**B**). Confocal image closeup of pereopod three carpus silk glands, with smaller oval epicuticle glands (**C**). Light microscope image of bromophenol blue stained merus of pereopod four (**D**). SEM of pereopod four dactylus tip (**E)**. SEM close up of pereopod four dactylus tip; small silk threads can be seen exiting the pores (**F**). Light microscope image of bromophenol blue stained pereopod four dactylus tip (**G**). Illustrated diagram of silk-spinning dactylus; setal grooves and pores (light grey), secretion points from pores (dark purple), gland ducts (light purple), dactylar channel (yellow), and setae (black) (**H**).

## DISCUSSION

The silk-spinning system in ampeliscoids is more similar to corophioids than previously thought, with a two-gland system distributed within the third and fourth pereopods, throughout the basis, ischium, merus, and carpus. Type one glands share similarities with the proximal glands found in corophioids, which have dense secretory products in the secretory cell (Nebeski 1880; Kronenberer *et al*., 2012; Neretin 2016; Neretin *et al*., 2017). Type two glands are similar to the distal glands of corophioids, which have an “empty” secretory cell with dense accumulation sites stained strongly with bromophenol blue (Nebeski 1880; Kronenberer *et al*., 2012; Neretin 2016; Neretin *et al*., 2017). Both gland types could be considered pseudotubular, as seen in corophioid silk glands (Neretin *et al*., 2017). The secretory products in type two glands were weakly stained by bromophenol blue stain, which targets acidic proteinaceous products, providing additional evidence that these glands are likely producing different products than gland type one. *Ampelisca* species contained type one glands with varying secretory cells, which we dubbed gland types 1A and 1B (Fig. 2F). We believe that this variation in staining is due to differences in the secretory products of the gland rather than variation in staining efficacy, likely indicating that multiple types of products can be produced from a single gland. This is similar to variation seen in the proximal glands of *Ampithoidae* species (Coriophidae; Neretin 2016) and silk glands of some genera of spiders (Sonavane *et al*., 2023)

The silk extrusion morphology of the dactyl tips was also more similar between ampeliscoids and corophioids than expected. Most prominently, a reservoir chamber was found in ampeliscoids. This chamber is suspected to play a role in silk storage (Kronenberger et al., 2012) and the mixing of products from both gland types (Neretin 2016). It has only been described in corophioids, and is known from no other silk-producing arthropods. We found that ampeliscoids have a single duct connecting to the reservoir chamber and did not find evidence of both gland types mixing before or within the chamber. Other ducts appear to have their own pores. This suggests that ampeliscoids are able to produce silk threads without the combination of secretory products and that mixing products is not a requirement for amphipod silk. It may be that ampeliscoids still use the chamber for storage, but it does not assist in mixing products. Ampeliscoids may transform silk more similarly to other arthropods such as silk moths (*Bombyx mori* Linneaus, 1758) utilizing an elongated duct and thickened duct. Silk moths have an elongated duct followed by a thickened cuticular duct wall, called a cuticular intima, before the exit pore (Huang *et al*., 2025). The cuticular intima applies shear force to the liquid silk which forces molecules into crystalline configurations, like β-sheets, creating a solid thread before exiting the body (Wang *et al*., 2022; Huang *et al*., 2025). The ampeliscoids silk extrusion morphology is similar, suggesting the elongated duct and thickened wall of the chamber may function similarly. In some corophioid species, an additional duct adjacent to the sensillum groove has been identified (Kronenberger *et al*., 2012). This duct may be analogous to the pores distal to the dactylar channel in ampeliscoids: the proximal pore in *Haploops*, the large terminal pore in *Ampelisca*, and second large terminal pore in *Byblis* (Fig. 3). Similar to corophioids, the main thread-producing pores appear to be on the opposite side of the dactyl tip to the setal groove. An additional distinction in ampeliscoids is the number of pores on the dactyl tip. In corophioids, only two pores are generally found. In contrast, *Ampelisca* can have an additional 16 small pores, *Haploops* six additional small pores, and *Byblis* one additional large pore. Based on ancestral state reconstruction, it appears that multiple pores is a derived trait that evolved in the common ancestor of *Ampelisca* and *Haploops* (Fig. 5) and is not ubiquitous in ampeliscoids.

**Figure 5.**
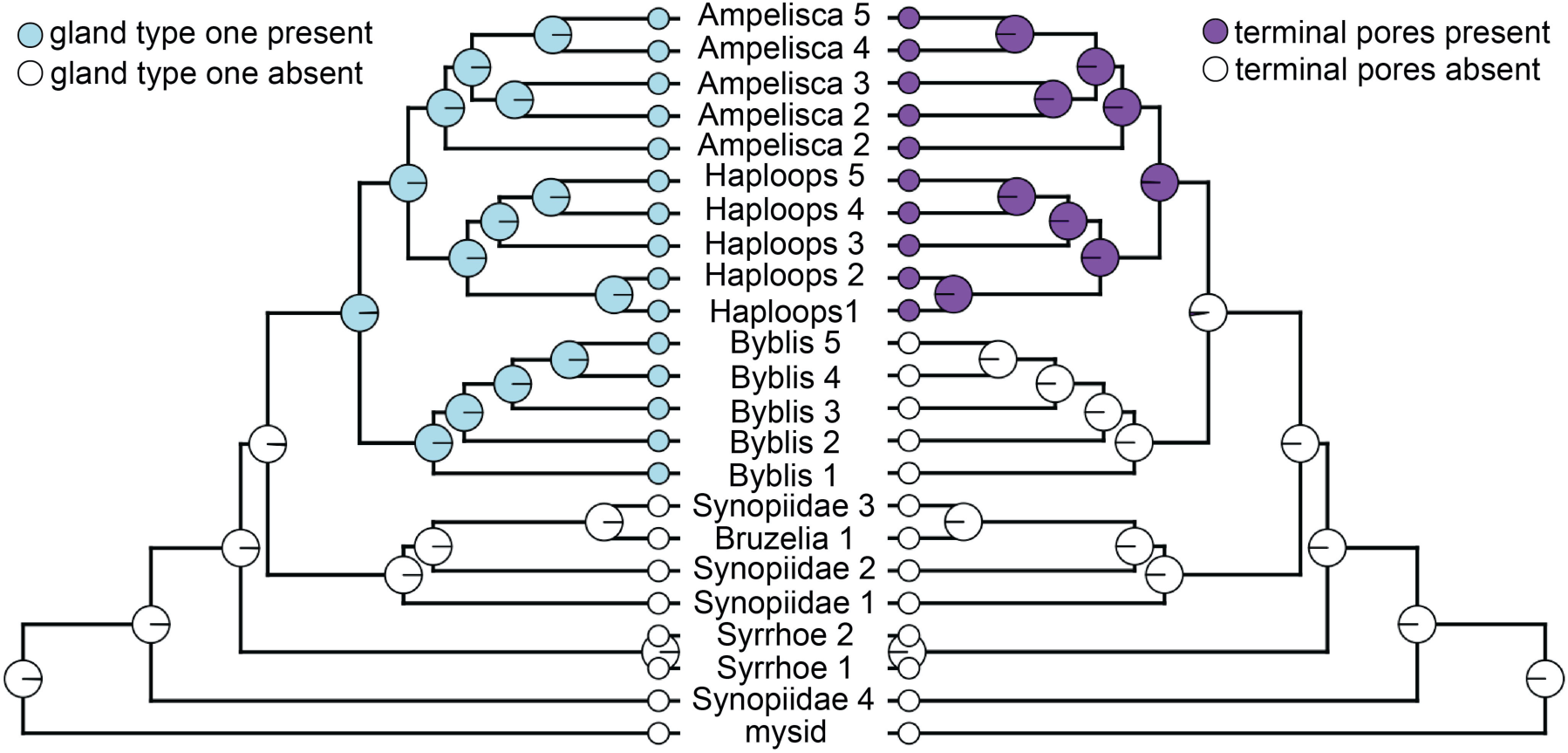
Ancestral State Reconstruction of key silk-spinning morphology; gland type one and dactylar terminal pores. Pie charts at nodes indicate the marginal probability of the trait present or absent in the common ancestor.

Similar to corophioid silk domiciles, both ampeliscoid and paragammaropsid domiciles were found to be composed of silk thread of varying thickness, mixed with amorphous secretions (Fig. 6). This supports the claim that these amphipods can produce a diversity of secretions, with some products having more specialized functions.

**Figure 6.**
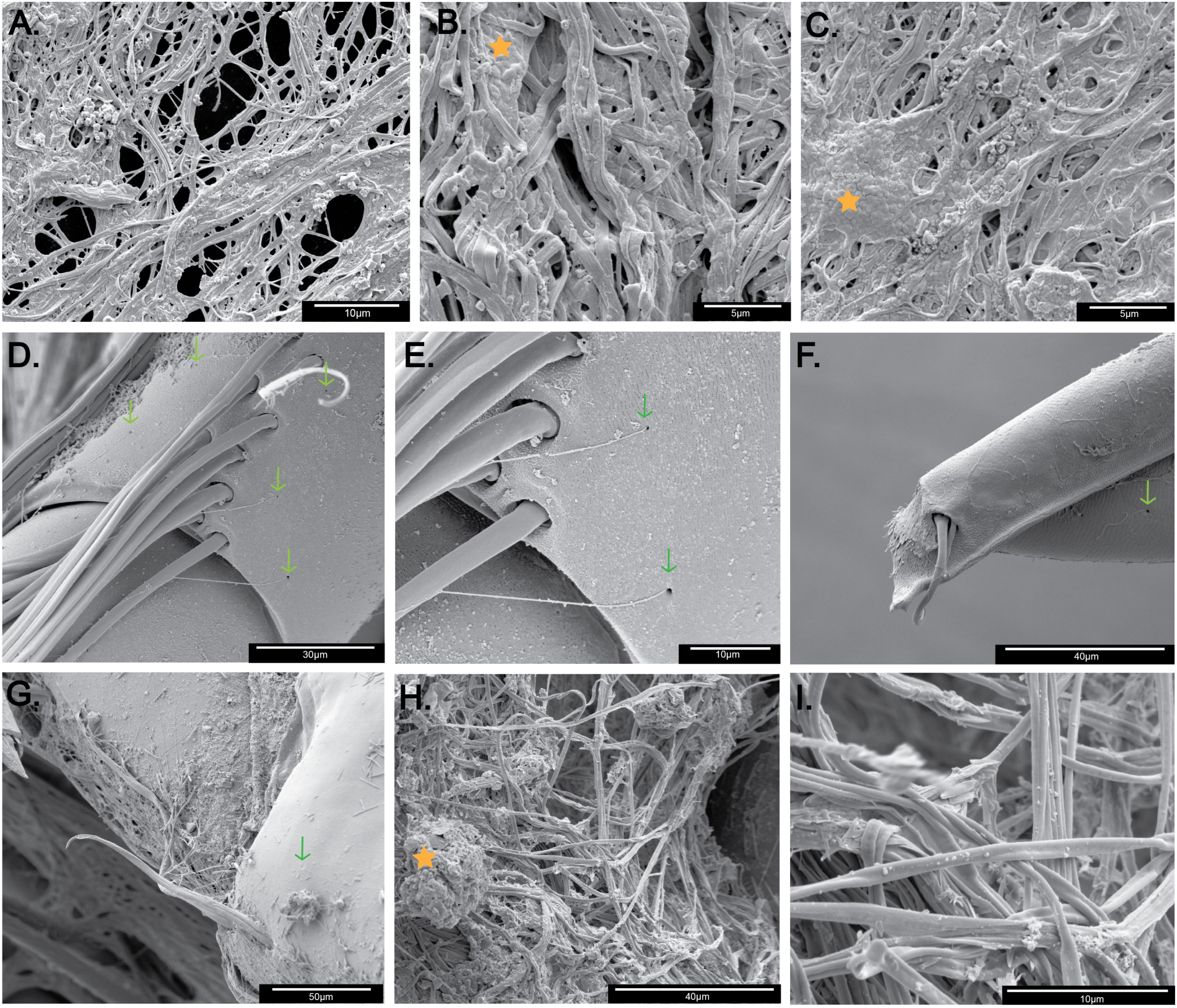
Spun silk and cuticle silk pores. Amorphous secretions (orange star) and cuticle pores (green arrows). A-F from *A. lobata*. Spun silk from domicile (**A**). Close up of silk (**B**). Close up of silk (**C**). Cuticle pores on pereopod three carpus with thin threads exiting (**D**). Close up of cuticle pores with threads (**E**). Pereopod seven dactylus with cuticle pore and thin thread exiting (**F**). *H. setosa* pereopod four distal merus with cuticle pore (**G**). *Paragammaropsis sp.* spun silk from domicile (**H**). Close up of spun silk (**I**).

Other pereopods of corophioids have been observed producing threads (Mattson & Cedhagen 1989), but little is known about their associated glands. This makes it difficult to compare pereopods two, five, six, and seven, between ampeliscoids and corophioids. In ampeliscoids we found both gland types in all examined pereopods (pereopods two to seven). It appears that these glands can produce thin threads from pores on the surface of the cuticle (Fig. 6D-G) but likely do not produce the thicker silk threads found on the domiciles (Fig. 6A-C). These cuticle pores may be analogous to the “muciferous” pores found on the cuticle of corophioids (Mattson & Cedhagen 1989). Further analyses are needed to confirm which gland types are associated with these cuticle pores. Though we don’t yet know if these products are proteinaceous or mucus, these findings further support the argument that mixing gland products are not required for amphipods to produce threads. It has been observed that ampeliscoids use secretions form pereopods one and two to construct a framework for their sandy domiciles and finish the final structure with pereopod three and four secretions (Mills 1967). These thin threads may play a role in preparing sediment for domicile construction, creating a framework for the domicile, or to repair the domicile, similar to corophioids (Shillaker & Moore 1978; Mattson & Cedhagen 1989). Ampeliscoids orient themselves with their head and posterior pereopods at the entrance to their domicile (Segerstråle *et al*., 1949). If they are producing thin threads to repair the domicile entrance, this could explain why there may be a higher abundance of thread pores on pereopod seven. In *Haploops* species, we found large type two glands in pereopod two (Fig. 2H), similar to those found in a previous study (Rigolet el al., 2011), that were larger and more abundant than seen in other genera. *Haploops* species are strict suspension feeders, unlike *Ampelisca* and *Byblis* species which do both suspension feeding and deposit feeding (Enequist 1949; Esposito *et al*., 2014), which may explain this difference. To help suspension feed, *Haploops* species rub their second pereopods on their antennae, transferring adhesive secretions which can be used to capture food particles in the water (Rigolet *et al*., 2011).

The presence of gland type two in synopiids suggests a potential origin of silk-spinning glands in ampeliscoids. We found type two glands in all examined pereopods of synopiids but found no evidence of silk threads being produced externally. Considering that type two glands were likely present in the common ancestor of ampeliscoids and synopiids (Fig. 5), we hypothesize that type two glands are ancestral glands that may be the precursor to type one glands. With the origin of type one glands, the silk extrusion morphology in pereopods three and four evolved as well, allowing the production of durable silk threads used for domicile building. Because similar glands and silk extrusion morphology can be found in corophioids, we hypothesize that type two glands are ancestral silk glands for both groups, and that type one glands evolved from a subfunctionalization event of ancestral type one glands. Figure 7 illustrates this hypothesis further. Additional morphological studies on the glands in corophioid pereopods two and five to seven, and in outgroup species, would help to advance this hypothesis. Although corophioids were not included in this analysis, we expect outgroups of corophioids to have type two glands as well, especially since pereopodal glands may be more prevalent in amphipods than previously known (based on our own preliminary, unpublished data). In many cases, it appears that silk glands across arthropods evolve from preexisting glands with a different ancestral function and then secretory cells of the gland undergo subfunctionalization to develop silk production. For example, this process has been found in the malpighian tubules and labial glands of insects, where excretory cells undergo subfunctionalization to produce silk (Spiegler 1962; Sutherland *et al*., 2010).

**Figure 7.**
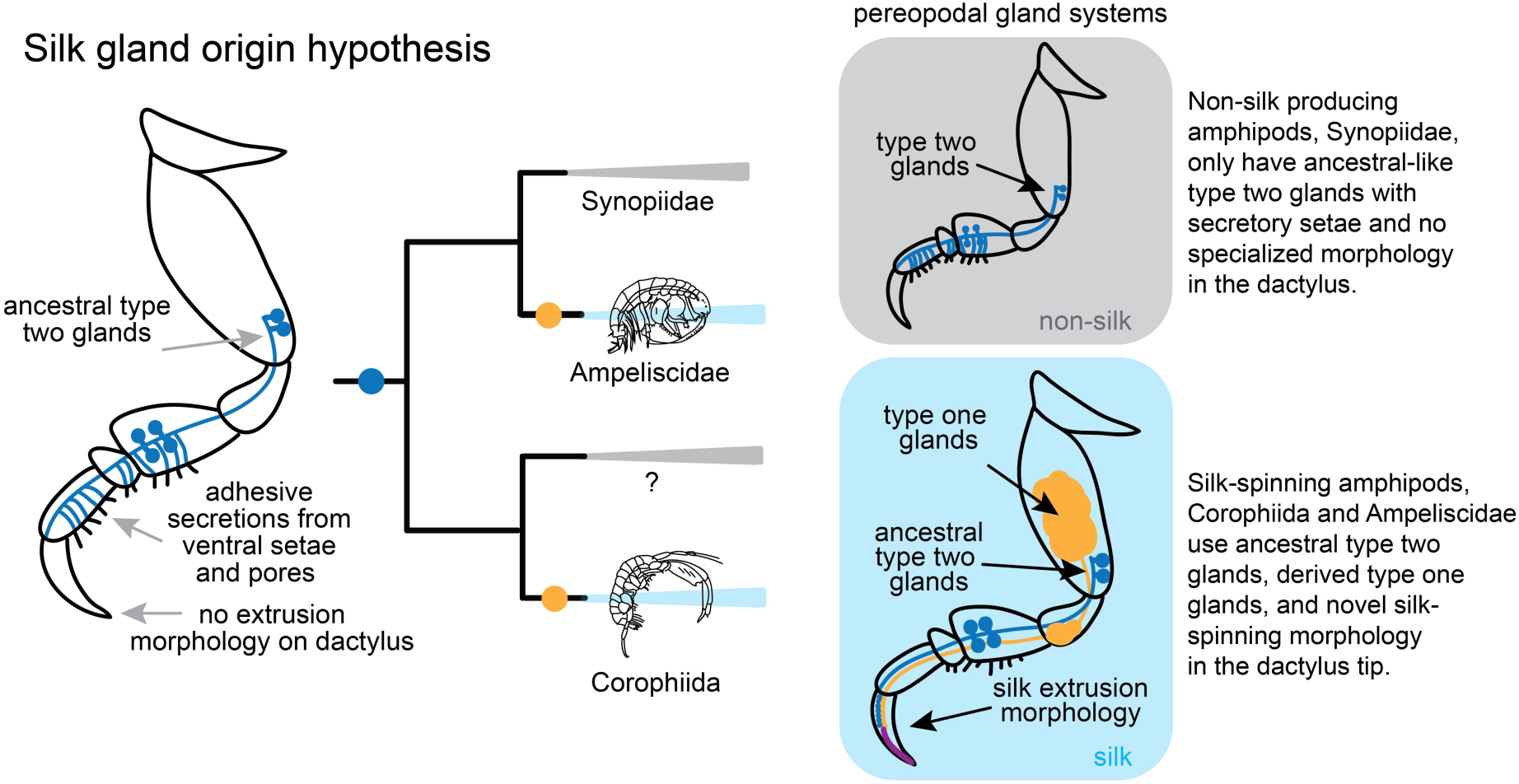
Hypothesis on the origin of silk-spinning glands in Senticaudata amphipods.

The overlap in silk-spinning systems in ampeliscoids and corophioids raises questions about the relationship between these two groups. Are these two groups sister clades, sharing silk glands that evolved in a common ancestor? Or are there developmental constraints or selective pressures within amphipods that facilitate strikingly convergent silk-spinning systems? If type two glands are preadaptations of silk-spinning systems in these two groups, similar constraints and pathways may apply to both lineages essentially evolving in parallel. Building robust phylogenies from genome-wide and transcriptomic data are therefore a high priority for future work.

Previous work (Myers and Lowry 2003) used the glandular distribution in pereopods three and four as a taxonomic trait to inform amphipod classification. Based on this system, if the glands are limited to the basis, this supports classification as Corophiida, whereas glands limited to the merus support classification as Ampeliscidae. Recent literature, however, has found glands beyond the basis in corophioids (Kronenberger *et al*., 2012; Neretin 2016; Neretin *et al*., 2017). Our data confirm and extend these findings in ampeliscoids, with silk-spinning glands that are distributed throughout the entire pereopod. Therefore, this trait should be reconsidered for classifying these taxonomic groups. Other work using phylogenetic estimations has suggested that the parvorder Synopiidira (synopiids) and ampeliscoids should be placed within the suborder Senticaudata, which also contains the Corophiida (Copilaş-Ciocianu 2020). If this is the case, then this presents a scenario in which type two glands may be synapomorphy for Senticaudata and supports the argument that type one glands are derived. Moores and Myer 1998 additionally suggested that if glandular pereopods are homologous in amphipods, then Ampeliscidae and Corophiida are likely to be sister taxa.

The Paragammaropsidae are an interesting group relevant to these considerations. *Paragammaropsis* has type one-like glands restricted to pereopod three and four, in contrast to ampeliscoids where they are present in pereopods two to seven (Fig. 4A-D). The species we examined, however, lacked type two glands in these pereopods, and instead had type two glands only in pereopod six. In this case, silk-spinning from pereopods three and four only relies on gland type one which is dissimilar to both groups. The silk extrusion morphology of *Paragammaropsis* shares similarities with ampeliscoids by having many small terminal pores, a reduced reservoir chamber like *Ampelisca,* and a dactyl channel running laterally from one setal groove to the other like *Byblis* (Fig. 4E-H). From these pores, and the terminal pore, *Paragammaropsis* can produce an abundance of fine silk threads (see Fig. 4F for silk coming out of pores, and Fig. 6H & I for spun silk). Based on glandular morphology alone, *Paragammaropsis* is therefore more similar to ampeliscoids, but these data alone seem insufficient to confidently place the Paragammaropsidae. Further phylogenetic analyses are suggested for a more conclusive answer.

Our findings provide the first clear evidence that the silk-spinning pereopods of ampeliscoids closely parallel those of corophioids, which motivates a re-evaluation of their taxonomic and phylogenetic relationship. At the same time, key differences in ampeliscoid silk-spinning systems, particularly their ability to produce threads without combining multiple gland products, reveal that amphipod silk production is more evolutionarily versatile than previously recognized. Building on these insights, we propose a new hypothesis for the origins of silk spinning in both groups, underscoring the broader significance of investing underexplored amphipod lineages. Expanding morphological studies across these taxa will be essential for unraveling the evolution of complex glandular systems and their ecological roles. Future work should prioritize additional ampeliscoid diversity, especially the elusive *Byblisoides* Barnard 1931, as well as a wider range of corophioid outgroups, to further refine and test this emerging evolutionary framework. This work establishes an essential comparative foundation for investigating how silk production has independently diversified across arthropod lineages.

## Supporting information

S1 Table; S2 Table; S3 Protocol

## SUPPLEMENTARY MATERIAL

S1 Protocol. Bromophenol blue stain recipe and protocol

S2 Table. Samples used for morphological investigation

S3 Table. cox1 gene accession numbers from NCBI used for phylogenetic inference 3

This project would not be possible without our funding support: The NSF-GRFP (22-614), The UCSB Coastal Fund Grant (CF-202511-15626), and The Grants in Aid of Research from Sigma Xi (G20250315-12637).

## ACKNOWLEDGMENTS

Thank you anonymous reviewers for your feedback. Thank you to Skylah Reis and Gustav Paulay for the collection of specimens in Santa Barbara, Kevin Kocot for collecting and sharing specimens from Antarctica, and Adam Wall for loaning specimens from the Natural History Museum of Los Angeles. Thank you to Benjamin Lopez and Lisa Mesrop for helping to develop staining protocols and assisting with microscopy techniques. Thank you to Steffen Harzch’s lab at the University of Greifswald Germany (Steffen Harzch, Frederice Hilgendorf, Alexandre Casadei-Ferreira, Sebastian Büsse, and Jan Phillipp Geißel) for preparing specimens for micro-CT and for providing the assistance, training, and equipment needed for this analysis. Thank you to Jacob Gorneau for helping with phylogenetic estimation methods. Thank you to Lewis "Lee" Sharpnack and Gareth Seward at the UCSB Earth Science Electron Microscopy and Microanalysis Lab for their assistance with SEM data collection. We acknowledge the use of the NRI-MCDB Microscopy Facility and the Resonant Scanning Confocal supported by the NSF MRI grant DBI-1625770.

## FIGURE ALT TEXT

Figure 1. **ALT TEXT:** An evolutionary tree of Ampeliscidae amphipods and outgroups: Synopiidae species and a distant outgroup, arctic mysid. It shows the phylogenetic relationship of these groups. Ampeliscidae species are all together in one monophyletic group.

Figure 2. **ALT TEXT:** Illustrations and microscopic images labelled A to G. A is an illustration of a generic ampeliscoid amphipod showing where glands are found in the legs with the number of pereopod labelled. B. is an illustration of a generic pereopod three and four with glands distributed throughout the leg. C. is an illustration of a generic non-silk-spinning pereopod with glands distributed throughout the leg. D. has four microscopic images of the pereopodal glands stained in Byblis species. E. has four microscopic images of the pereopodal glands stained in *Haploops* species. F. has four microscopic images of the pereopodal glands stained in *Ampelisca* species. G. has four microscopic images of the pereopodal glands stained in synopiid species.

Figure 3. **ALT TEXT:** Illustrations and microscopic images labelled A to C showing the silk extrusion morphology of Ampeliscidae genera. A has SEM and light microscopy images, and a summary illustration of the dactylus tip of *Byblis* species highlighting three pores where silk exits. B. has SEM and light microscopy images, and a summary illustration of the dactylus tip of *Haploops* species highlighting two main pores and many small pores where silk exits. C. has SEM and light microscopy images, and a summary illustration of the dactylus tip of *Ampelisca* species highlighting two main pores and many small pores where silk exits.

Figure 4. **ALT TEXT:** 3D volume, microscopic images, and an illustration of the silk-spinning system of *Paragammaropsis sp.* labelled A to H. A is a 3D volume of pereopod four showing large silk glands in the merus and carpus. B is a confocal image of pereopod four with glands in the merus and carpus. C is a confocal image closeup of pereopod three carpus filled with glands. D is a light microscope image pereopod four merus stained with bromophenol blue. E is a SEM of pereopod four dactylus tip. F is a SEM closeup of pereopod four dactylus tip showing small terminal pores and larger terminal pore. G is a light microscope image of pereopod four dactylus tip. H is an illustration summarizing the key features of the dactylus tip with many small pores and a larger pore.

Figure 5. **ALT TEXT:** Evolutionary tree of Ampeliscidae and outgroups with morphological character states at the tips and at the internal nodes with pie charts representing the marginal probability of the trait present or absent in the ancestor. Left tree with gland type one character and right tree with dactylar terminal pores character.

Figure 6. **ALT TEXT:** SEM images of the silk from *A. lobata, H. setosa,* and *Pargammaropsis sp.* Amphipods, labelled A to I. A is spun silk from the domicile of *A. lobata*. B is a close up of the spun silk. C is another close up of the spun silk. D is the cuticle pores on pereopod three carpus of *A. lobata*, with threads coming out. E is a close up of those pores. F is the cuticle pore on pereopod seven of *A. lobata* with a thread coming out. G is the cuticle pores and silk from pereopod four merus of *H. setosa*. H. is spun silk from the domicile of *Paragammaropsis sp.* I is a close up of the spun silk.

Figure 7. **ALT TEXT:** An illustration of a simplified evolutionary tree with silk-spinning amphipods from Ampeliscidae and Corophiida. The figure shows the silk-spinning glands found in modern taxa and a hypothetical drawing of what the ancestral glandular system might have looked like.

## Notes

### Competing Interest Statement

The authors have declared no competing interest.

