## Supplementary material for "Comparative morphology of silk-spinning systems in amphipods": S1 Table; S2 Table; S3 Protocol

cox1 genes from NCBI used for phylogenetic inference

| Name on ASR tree | species name | accession number NCBI |
|---|---|---|
| Haploops1 | Haploops_robusta | OQ053036.1 |
| Haploops2 | Haploops_carinata | OQ053034.1 |
| Haploops3 | Haploops_antarctica | PQ846648.1 |
| Haploops4 | Haploops_tubicola | MG934885.1 |
| Haploops5 | Haploops_tenuis | PP192150.1 |
| Ampelisca1 | Ampelisca_macrocephala | PV055614.1 |
| Ampelisca2 | Ampelisca_brevicornis | JQ653148.1 |
| Ampelisca3 | Ampelisca_eschrichtii | HM425345.1 |
| Ampelisca4 | Ampelisca_typica | JQ653138.1 |
| Ampelisca5 | Ampelisca_diadema | JX100839.1 |
| Byblis1 | Byblis_sp. | PV077105.1 |
| Byblis2 | Byblis_crassicornis | OQ053048.1 |
| Byblis3 | Byblis_veleronis | HQ941723.1 |
| Byblis4 | Byblis_gaimardi | DQ889084.1 |
| Byblis5 | Byblis_frigidis | MG320512.1 |
| Synopiidae1 | Syrrhoites_sp. | MG264811.1 |
| Synopiidae2 | Syrrhoites_pusilla | MG264798.1 |
| Synopiidae3 | Synopiidae_sp. | MN346625.1 |
| Synopiidae4 | Synopia_sp. | EF989701.1 |
| Syrrhoe1 | Syrrhoe_crenulata | MK972328.1 |
| Syrrhoe2 | Syrrhoe_sp. | MG319247.1 |
| Bruzelia1 | Bruzelia_sp. | MG264865.1 |
| Mysid | Antarctomysis_maxima | PQ846639.1 |

**S3 Table**

Samples used for morphological investigation

| species | Experiment_ID | Loan source | Loan_Collections_ID |
|---|---|---|---|
| Ampelisca_lobata | SM667A | collected by Siena Mckim | NA |
| Ampelisca_lobata | SM92 | collected by Siena Mckim | NA |
| Ampelisca_lobata | SM667B | collected by Siena Mckim | NA |
| Ampelisca_romigi | SM585A | Natural History Museum of Los Angeles County | 253740 |
| Ampelisca_romigi | SM586A | Natural History Museum of Los Angeles County | 240850 |
| Ampelisca_pacifica | SM587A | Natural History Museum of Los Angeles County | 274308 |
| Ampelisca_pacifica | SM588A | Natural History Museum of Los Angeles County | 275778 |
| Byblis_veleronis | SM342A | Natural History Museum of Los Angeles County | 217456 |
| Byblis_veleronis | SM342B | Natural History Museum of Los Angeles County | 217456 |
| Byblis_millsi | SM589A | Natural History Museum of Los Angeles County | 255392 |
| Byblis_millsi | SM590A | Natural History Museum of Los Angeles County | 244809 |
| Haploops_laevis | SM637 | Natural History Museum of Los Angeles County | 411454 |
| Haploops_setosa | SM639A | Natural History Museum of Los Angeles County | 286390 |
| Haploops_setosa | SM640 | Natural History Museum of Los Angeles County | 422421 |
| Paragammaropsis_sp. | SM469A.1 | Alabama Museum of Natural History | ALMNH:Inv:21923 |

| | | | |
|---|---|---|---|
| Paragammaropsis_sp. | SM469A.2 | Alabama Museum of Natural History | ALMNH:Inv:21923 |
| Paragammaropsis_sp. | SM469A.3 | Alabama Museum of Natural History | ALMNH:Inv:21923 |
| Bruzelia_tuberculata | SM606 | Natural History Museum of Los Angeles County | 469992 |
| Bruzelia_tuberculata | SM607A | Natural History Museum of Los Angeles County | 469272 |
| Tiron_biocellatus | SM608 | Natural History Museum of Los Angeles County | 262153 |
| Tiron_biocellatus | SM610A | Natural History Museum of Los Angeles County | 293852 |
| Syrrhoe_longifrons | SM533A | Santa Barbara Museum of Natural History | 460150 |

**S3 Protocol**

Bromophenol Blue Stain Recipe and Procedure

**Mercury bromophenol blue recipe - makes 50mL**

50mL of 95% ethanol
50mg bromophenol blue powder stain
5g mercury chloride $HgCl_2$

**Bromophenol blue destainer - makes 50mL**

.25mL of acetic acid
49.75 mL DI water

**Stain target**

Bromophenol blue is intended to stain sulphydryl, phenolic and carboxyl groups of proteins.

**Procedure**

If starting with specimens preserved in 75% ethanol, incubate specimens in 80% ethanol for 5 minutes and then in 100% ethanol for 5 minutes to dehydrate the specimen. Then place it into a well of mercury bromophenol blue for 3-30 minutes depending on the thickness of the specimen. Do not wait until the specimen is entirely blue, it will continue to darken after being taken out. Next, de-stain background stain with .5% acetic acid wash for 20 minutes. Next, place specimens in tert-butanol ($C_4H_{10}O$) that has been warmed in a hot water bath. Rinse 3 times, each time for 5 minutes, then incubate for 1.5 hours and complete one more wash. Incubate for a final 1.5 hours. For each rinse, make sure tert-butanol has been warmed, or else it may crystallize and damage specimens. And keep specimens incubating on a hot water bath, ensuring no water splashes into the vessel. Lastly, place the specimen into a well of Histo-Clear for 1-5 hours depending on the thickness of the specimen cuticle. From here, you can mount your specimen onto a slide in 100% glycerol or you can store the specimen in a tube of glycerol in a 4°F fridge for up to 2 years (or longer depending on the preservation method of the specimen).

Recipe and procedure adapted from Bonhang 1955 and Kronenberger et al., 2012 after fine tuning with amphipod specimens stored in ethanol and formaldehyde.
